# Reprogramming the endogenous type III-A CRISPR-Cas system for genome editing, RNA interference and CRISPRi screening in *Mycobacterium tuberculosis*

**DOI:** 10.1101/2020.03.09.983494

**Authors:** Khaista Rahman, Muhammad Jamal, Xi Chen, Wei Zhou, Bin Yang, Yanyan Zou, Weize Xu, Yingying Lei, Chengchao Wu, Xiaojian Cao, Rohit Tyagi, Muhammad Ahsan Naeem, Da Lin, Zeshan Habib, Nan Peng, Zhen F. Fu, Gang Cao

**Author notes:** equally contributed to this work as a first author (Khaista Rahman, Muhammad Jamal).

## Abstract

*Mycobacterium tuberculosis* (*M.tb*) causes the current leading infectious disease. Examination of the functional genomics of *M.tb* and development of drugs and vaccines are hampered by the complicated and time-consuming genetic manipulation techniques for *M.tb.* Here, we reprogrammed *M.tb* endogenous type III-A CRISPR-Cas10 system for simple and efficient gene editing, RNA interference and screening *via* simple delivery of a plasmid harboring a mini-CRISPR array, thereby avoiding the introduction of exogenous proteins and minimizing proteotoxicity. We demonstrated that *M.tb* genes were efficiently and specifically knocked-in/out by this system, which was confirmed by whole-genome sequencing. This system was further employed for single and simultaneous multiple-gene RNA interference. Moreover, we successfully applied this system for genome-wide CRISPR interference screening to identify the *in-vitro* and intracellular growth-regulating genes. This system can be extensively used to explore the functional genomics of *M.tb* and facilitate the development of new anti-*Mycobacterial* drugs and vaccines.

**Summary:** Tuberculosis caused by *Mycobacterium tuberculosis* (*M.tb*) is the current leading infectious disease affecting more than ten million people annually. To dissect the functional genomics and understand its virulence, persistence, and antibiotics resistance, a powerful genome editing tool and high-throughput screening methods are desperately wanted. Our study developed an efficient and a robust tool for genome editing and RNA interference in *M.tb* using its endogenous CRISPR cas10 system. Moreover, the system has been successfully applied for genome-wide CRISPR interference screening. This tool could be employed to explore the functional genomics of *M.tb* and facilitate the development of anti-*M.tb* drugs and vaccines.

## Introduction

*Mycobacterium tuberculosis* (*M.tb*) is the powerful etiological agent of tuberculosis (TB). Currently, it is the deadliest pathogen, ranking above HIV, causing approximately 1.3 million deaths and 10 million new cases globally (Floyd et al., 2018). Currently, the treatment for TB comprises the extensive administration of antibiotics, which often leads to the emergence of extensively drug resistance bacteria (Mittal and Gupta, 2011). Meanwhile, the intake of rifampicin, isoniazid or pyrazinamide could adversely affect the composition of gut microbiota (Hu et al., 2019; Khan et al., 2019). Functional genomic analysis of *M.tb* and elucidation of the molecular mechanism underlying TB pathophysiology are crucial for identification of new drug targets and vaccine candidate genes. However, advances in *M.tb* functional genomic studies are greatly impeded by inefficient tools for gene editing and silencing. The conventional methods for gene editing based on simple homologous recombination using tools such as non-replicating plasmids (Husson et al., 1990), incompatible plasmids (Pashley et al., 2003), linear DNA substrates (Balasubramanian et al., 1996) and phage-based transduction (Bardarov et al., 2002) are less efficient. Although phage transduction can increase the efficiency of *M.tb* genetic manipulation, this technique is laborious and time consuming (Choudhary et al., 2016; Tufariello et al., 2014; Van Kessel and Hatfull, 2007). Recently a method called ORBIT (oligonucleotide-mediated recombineering followed by Bxb1 integrase targeting) was developed for gene editing in *M.tb* based on the integration of a targeting oligonucleotide in the genome via homologous recombination mediated by the phage Che9c RecT annealase. However, this method needs the transformation of a single stranded DNA probe along with a non-replicating plasmid (Murphy et al., 2018).

The clustered regularly interspaced short palindromic repeats (CRISPR) and associated genes (Cas) system has been extensively used for genome editing in bacteria (Jiang et al., 2013; Nishimasu et al., 2014). The CRISPR-Cas9 systems from *Streptococcus pyogenes* and *Streptococcus thermophilus* have been used for sequence-specific transcriptional repression in *M.tb*, achieving substantial gene silencing (Choudhary et al., 2015; Rock et al., 2017; Singh et al., 2016). Recently, Rock. et al screened a group of Cas9 proteins for gene silencing in *M.tb* and found that dCas9 from *S. thermophiles* was more efficient for gene silencing in *M.tb* with a reduced proteotoxicity (Rock et al., 2017). Yet, the application of CRISPR-Cas9 failed to disrupt genes in *M.bovis, M.smegmatis* and *M.bovis* BCG. (Sun et al., 2018; Vandewalle, 2015). Moreover, to date, none of these methods have been applied to simultaneously silence multiple genes to dissect gene function and study genetic interactions in *M.tb*.

The type III CRISPR-Cas system is present in approximately 75% of archaea and 40% of bacteria, including pathogenic *Mycobacterium* and *Staphylococcus* species (Makarova et al., 2011). This system is further classified into four subtypes that are characterized by the Csm complex (type III-A and D) or Cmr (type III-B and C) (Koonin et al., 2017). The initial transcription of CRISPR array yields an immature long transcript known as pre-crRNA, which is processed by Cas6 along with other nuclease to produce mature crRNAs. The mature crRNA harbors an 8nt tag at its 5’-end known as “crRNA-tag”, which is of pivotal importance in auto-immunity scenario. (Carte et al., 2008; Deltcheva et al., 2011; Hatoum-Aslan et al., 2011; Estrella et al., 2016; Hatoum-Aslan et al., 2011; Kazlauskiene et al., 2016). Once the mature crRNA has been generated, the Csm protein complex interacts with the crRNA to form a ribonucleoprotein complex (Samai et al., 2015).

The type III CRISPR-Cas system possess transcription dependent RNA and DNA cleavage capabilities, hence efficiently providing immunity against invading genomic elements in *Staphylococcus epidermidis* (Koonin and Makarova, 2013; Samai et al., 2015; Guo et al., 2019). The RNA cleavage is mediated by the Csm3/cmr4 or csm6/csx1 complex, while the DNA cleavage is catalyzed by Cas10 (Csm1) (Grüschow et al., 2019a; Park et al., 2017; Samai et al., 2015; Tamulaitis et al., 2017).). Upon activation, Cas10 cleaves the ssDNA (Estrella et al., 2016; Kazlauskiene et al., 2016) and generates the cyclic oligoadenylate (cOA) signaling molecules from ATP, which further act as an activator for the RNA targeting proteins (Kazlauskiene et al., 2017; Niewoehner et al., 2017).

The type III CRISPR system can be reprogrammed for RNA editing in *Streptococcus thermophiles* and *Sulfolobus islandicus* (Tamulaitis et al., 2014 Li et al., 2015; Peng et al., 2015). Meanwhile, this system has also been reprogrammed for chromosomal targeting and to achieve large-scale genomic deletion and alteration in *Staphylococcus aureus* (Guan et al., 2017). Bari et al. has also been used this system for DNA editing of Virulent *Staphylococcal* phages (Bari et al., 2017). Recently, Wei et al. reported that *Mycobacterium* species contain a typical features of type III-A systems and are highly active (Wei et al.). More recently Grüschow et al. showed that the *M.tb* CRISPR system effectively targets RNA and DNA upon expression in *E.coli* (Grüschow et al., 2019b; Wei et al., 2019). However, application of this system in *M.tb* for gene editing, RNA interference and CRISPR interference (CRISPRi) screening has not been reported.

The aim of this study is, therefore, employing the *M.tb* endogenous type III-A CRISPR-Cas system to develop a versatile and robust tool for efficient gene editing, RNA interference and genome wide CRISPRi screening. We demonstrate that this system can be used for robust and facile gene knock-in/out. Moreover, this system can be utilized for single and simultaneous multiple-gene RNA interference to precisely dissect the functions of specific genes. Furthermore, we applied this system for genome-scale CRISPRi screening of growth-regulating genes. This “killing two birds with one stone” tool may shed light on genetic and functional studies of *M.tb* and facilitate the development of anti-*M.tb* drugs and potent vaccines.

## MATERIALS AND METHODS

### Bacterial culture, competence cell preparation, eletrotransformation and *in-vitro* growth analysis

For culturing *Escherichia coli* (*E.coli*) the bacteria was cultured in lysogeny broth (LB) medium (with or without kenamycin) for 12-14 hours at 37 ºC in a shaking incubator. For culturing the *M.tb*, the baccili was cultured in Middle brook 7H9 broth supplemented with (Becton Dickinson) supplemented with 10 % OADC (oleic albumin dextrose catalase), 0.02 % tween 80 and 0.5 % glycerol or on Middle brook 7H10 agar petri plate supplemented with 10% OADC and 0.5 % glycerol containing kanamycin (25 μg/mL) when needed. For *in-vitro* growth analysis, the bacteria were cultured in Middle brook 7H9 broth and adjusted OD_600nm_ to 0.15. Bacterial suspension (0.2 mL) was used to examine OD_600nm_ each day for 18 days. All the experiments were performed in triplicates for each sample.

The competence cells were made with the following manner: 100 mL of *M.tb* H37Ra was pelleted by centrifuging at 8,000g for 10 minutes, 4 ºC and washed 3 times with 10 % ice chilled glycerol. The pellet was washed with 4 mL of 10 % ice chilled glycerol three times and aliquot into 80 Eppendorf tubes with 200 μL suspension. For electro-transformation, *M.tb* H37Ra was electro-transformed using Gene Pulser Xcell (BIO-RAD) with the default setting of 2.5 kV, 25 µF, and 1000 Ω resistance. The bacterial suspension was transferred to a fresh Eppendorf tube containing 1 mL of Middle brook 7H9 broth. The cell suspension was pelleted after 24 h incubation and plated onto Middle brook 7H10 agar plates containing kanamycin.

### Cell culture and infection

Human monocyte cell line (THP1, ATCC^®^ TIB-202) was cultured in RPMI-1640 complete growth medium supplemented with 10% FBS, and were differentiated for 24 h using culture medium containing 40 ng/mL phorbol 12-myristate 13-acetate (PMA) before infection. For infection, the cells were seeded in a T75 flasks for 24 h and were incubated with H37Ra bacilli (MOI=10) for 6 h at 37°C in 5% CO_2_. The cells were washed three times with pre-warmed PBS to remove extracellular bacilli, and supplied with fresh RPMI complete medium containing amikacin (50 μg/mL). The intracellular bacteria were isolated 3 days post infection and subjected plasmid isolation.

### Plasmid construction and preparation

To generate pMV-261-crRNA, two 36 bp repeats from *M.tb* H37Ra type III-A CRISPR array flanked by *Bbs*I sites on both side commercially were synthesized (Wuhan GeneCreate Biological Engineering Co., Ltd., Wuhan, China) and cloned into pMV-261. A 40bp guide RNA sequence corresponding to the sequence of the spacer in CRISPR-array was selected from the target genes. The spacer fragments were generated by annealing of the corresponding complementary oligonucleotides and cloned into the pMV-261-crRNA plasmid at *Bbs*I sites to generate pMV-261-gRNA. The guide RNA sequences corresponding to each gene are listed in Table S1. Homology-directed repair (HDR) templates (approximately 400bp) containing either GFP or BFP was generated by overlapping PCR using specific primers(Table S2) and inserted into pMV-261-crRNA at *Pst*I site. For gene knock-out study, the templates were generated by replacing the whole coding sequence of the target gene with GFP or BFP. To carryout gene interference, a 40 bp guide RNA sequence from the non-coding strands of the target gene flanked by the penta-nucleotide motif (5’-GAAAC-3’) of crRNA-tag (8 repeat handle) was taken as gRNA. The gRNA sequence of the genes and the controls are listed in Table S1. For multiple gene silencing, plasmid containing three spacers (40 bp) flanked by 36 bp repeats on both sides was commercially synthesized (Wuhan GeneCreate Biological Engineering Co., Ltd., Wuhan, China), of which the sequences are listed in Table S1.

### Fluorescence imaging

For transformants identification, the fluorescent colonies were either directly observed or re-suspended in sterilized PBS (Phosphate Buffer Saline) and spread on the microscopic slide, and then examined using Olympus IX73 microscope (Japan). Each selected colony was imaged for several different views using a filter with excitation wavelength of 465 - 495 nm and an emission filter of 515 - 555 nm for GFP and 55 - 375 nm and barrier filter of 400 nm for BFP. All Experiments were performed in triplicate.

### Identification of gene knock-in/out

Specific primers designed in the upstream and downstream of the HDR template or in the fluorescent gene shown in Table S2 were used for PCR screening of mutant strains. Forward primer (F) and reverse primer (R) designed upstream and downstream of the left arm and right arm of HDR template respectively, and R1 primer designed in e knock-in were used for PCR amplification (Figure S1. B). The PCR product was sequenced by Sanger sequencing (TsingKe Biological Technology, Wuhan, China) and analyzed by NCBI BLAST.

### RNA extraction and quantitative real-time PCR

*M.tb* H37Ra were pelleted and re-suspended in 1 mLTrizol Reagent (TAKARA), and then homogenized using Lysing Matrix B (MP Bio Medical, USA). 0.2 mL of chloroform was added to the bacterial suspension and centrifuge at 10,000 rpm for 10 min at 4 °C, of which the aqueous layer were transferred to a new RNase free Eppendorf tube. Isopropanol was used to precipitate RNA at room temperature for 10 min. RNA was then pelleted and washed twice with fresh made 80% EtOH. RNA pellet was air dried and dissolved in DEPC water and treated with TURBO DNase (Ambion) following manufacturer’s instruction to remove DNA. RNA was reverse transcribed to cDNA using ReverTra Ace^®^ qPCR RT Kit (Toyobo, Japan). The quantitative PCR was carried out using PerfeCTa SYBR Green SuperMix applied Biosystems 7300 real-time PCR system with Sequence Detection Software version v1.4.0. Data were analyzed using the 2^-ΔΔCT^ method with *gyrB* gene as the reference gene. The primers used for qRT-PCR are shown in Table S2.

### RNA immunoprecipitation q-PCR (RIP q-PCR) Assay

The RNA immunoprecipitation q-PCR (RIP q-PCR) assay was performed according to the method described by Minch, K. J. et al with some modifications (Minch et al., 2015). Briefly, bacteria was harvested from the mid-log phase and fixed with nuclease and proteases free 3% formaldehyde, lysed using Lysing Matrix B tubes followed by two rounds of bead beating at maximum speed for 40s. The tubes were then centrifuged at 4°C for 60s at 150g and the supernatant was transferred into a 0.5 mL Eppendorf tubes. The volume was normalized to 0.4 mL with CHIP buffer and sonicated. The samples were then incubated with 10 μg anti-HA CHIP grade antibody (Arigo bio lab, Cat. No ARG55095) at 4 °C for 8h. The csm complex was captured via HA-tagged csm6 by incubation with CHIP grade protein G magnetic beads. The tubes were then kept on a magnetic stand, the supernatant was removed and the beads were washed twice with CHIP buffer, twice with RNAse free IPP150 buffer and finally with 1X TE buffer. The RNA was extracted with Trizole method. After the DNA digestion, the mRNA was reverse transcribed into cDNA. The qRT-PCR was performed with *KatG* and *lpqE* specific primers (table S1). The data was normalized to the negative control and analyzed by using GraphPad prims 5.

### gRNA library designing and construction

The 40 base pairs sequence after the ‘GAAAC’ motif in *M.tb* H37Ra (Genome assembly: ASM1614v1) genome sequence was selected as the gRNA sequence candidates (Figure 7.1, S6 and Table S3). The library pool containing 5658 target fragments with identical ends were generated through on-chip oligo synthesis (Twist Biosciences USA) (Table S3). The oligo pool was dissolved in nuclease free TE Buffer (pH = 8.5) to a final concentration of 2.5 ng/mL and then was amplified with Phanta Max Super-Fidelity DNA Polymerase Kit (Vazyme biotech, China) using Lib-seq primer F and R (Table S2) for 12 cycles. The amplified product was run on 2% agarose gel and purified by OMEGA Gel purification Kit (OMEGA Bio-Tek). Next, the library was cloned into pMV-261-crRNA through homologues recombination using one step cloning kit (Vazyme biotech, China). The recombinant plasmids were then transformed into highly competent *E.coli* DH5α (Thremo fisher scientific catalogue #18265017). The efficiency and construction of library was validated by Sanger sequencing.

### Library screening and high throughput sequencing

For genome wide CRISPRi screening the plasmid DNA were isolated from all the three cultures of *M.tb* H37Ra. Then 20 µL from each plasmid was used as a template to amplify the crRNA along with repeats (Figure S4). 20 µL of the plasmid isolated from *E.coli* was also used as a reference. Briefly, 20 µL templates from each sample were amplified with library-seq primer F and R using Phanta Max Super-Fidelity DNA Polymerase Kit (Vazyme biotech, China). The PCR reaction was set as following: 98 °C for 4 min, 20 cycles (98 °C for 10 s; 56 °C for 10 s; 72 °C for 25 s), 72 °C for 5 min, 4 °C hold). The sequencing library of the gRNA library was prepared by VAHTS Universal DNA Library Prep Kit V3 for Illumina (Vazyme biotech, China) following the manufacturer’s instruction.

Briefly, the fragments were treated with End Prep Mix for end repairing, 5′ Phosphorylation, dA-tailing and purification using VAHTS DNA clean beads (Vazyme biotech, China). Then fragments were ligated with indexed adapters with a ‘T’ overhang. Subsequently, the ligated products were purified using the VAHTS DNA clean beads and amplified by PCR for 10 cycles. The libraries were run on gel, purified and validated by an Agilent 2100 Bioanalyzer (Agilent Technologies, Palo Alto, CA, USA). For *M.tb* H37Ra whole genome sequencing, the whole genomic DNA was isolated and subjected to fragmentation by sonication using Diagenode Bioruptor Pico System (Diagenode Inc. USA). The fragmented DNA was run on 1% agarose gel and the 150 to 350 bp fragments were obtained and purified, end repaired, ligated with adapters, and sequenced after amplification. Then the libraries with different indexes were pooled and loaded on Illumina X10 instrument for sequencing (Illumina, San Diego, CA, USA).

### High throughput sequencing data analysis

For *M.tb* whole-genome sequencing analysis, the adapters were trimmed from the raw data by Trimmomatic (Bolger et al., 2014). Clean data for each accession were mapped to the *M.tb* H37Ra reference genome (Genome assembly: ASM76770v1) using BWA (Li and Durbin, 2009) with default parameters. SNP and insertion-deletion (Indel) detection was achieved by using SAMtools and BCFtools (Danecek and McCarthy, 2017; Li et al., 2009; McKenna et al., 2010). The 20 bp up-stream and down-stream regions of the homologous segment in the guide RNA (gRNA) were selected as potential candidate off-target regions (Zumla et al., 2016). EDILIB was used (Sosic and Sikic, 2016)to align the gRNA against the reference genome with up to 15 mismatches, which was visualized as a bar plot. Bedtools (v2.26.0) was used to validate the gene knock-out (Quinlan and Hall, 2010).

For CRISPRi analysis, the raw data were first filtered according to the repeat-gRNA-repeat structure. Next, the gRNA sequences were extracted and counted. To normalize the total reads from different libraries to the equivalent size and maintain the read composition of each gRNA, the total count number of each sample was normalized using the following formula:

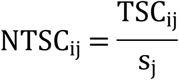

WhereTSC_ij_ represents the i^th^gRNA count in the samplej, and is the size factor of samplej. The median-of-ratios method in the DESeq R package was applied for this TSC normalization (Anders and Huber, 2010): 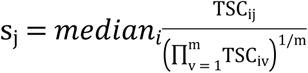

After normalization, the reads count ratio of each targets between *M.tb* and *E.coli* were calculated. For screening, the significantly increased/decreased targets, both *P*-value and the fold changes were considered. The independent *t*-test was performed to calculate the *P*-value and adjusted by BH method. *P*-value threshold is 0.05, fold change threshold is 4 for down counted targets, and 10 for up counted targets.

To check the similarity of significantly down counted *M.tb* proteins with the proteomes of the 43 most common probiotics (https://www.kegg.jp/kegg/catalog/org_list.html), the classic Needleman-Wunsch algorithm was applied (Needleman and Wunsch, 1970). The similarity is presented as a heatmap, which was generated using the pheatmap R package (https://cran.r-project.org/web/packages/pheatmap/).

## RESULTS

### Genomic and functional analysis of *Mycobacterium* type III-A CRISPR system

The CRISPR systems of three different species of *Mycobacterium* were explored and compared using the NCBI BLAST, Mycobrowser (https://mycobrowser.epfl.ch/), and KEGG genome databases (http://www.genome.jp/). The CRISPR systems of *M.tb* H37Rv and H37Ra comprise three CRISPR loci in association with 10 *cas* genes, namely, *cas2, cas1, csm6, csm5, csm4, csm3, csm2, cas10,cas6* and *cas4* (Figure 1.A). The CRISPR locus 1 (from left to right in Figure 1.A) of *M.tb* contains only two repeats flank by two spacers and is distant from the other two CRISPR loci and their associated *Cas* genes. Loci 2 and 3 are interspaced by two transposase genes belonging to the insertion sequence (*IS6110*) gene family followed by a cluster of 9 consecutive *cas* genes. Loci 2 and 3 contain 18 and 24 repeats, respectively, with a well-characterized leader sequence at the 5’ ends. Similarly, the CRISPR systems of *M.bovis* and *M.bovis* BCG have the same number and length of repeats and spacers as that of *M.tb* (Figure 1.A). However, the CRISPR system of *M.avium* has an uncharacterized and a fully characterized long CRISPR locus in association with 5 c*as4* genes and 11 repeats (Figure 1.A).

**Figure 1:**
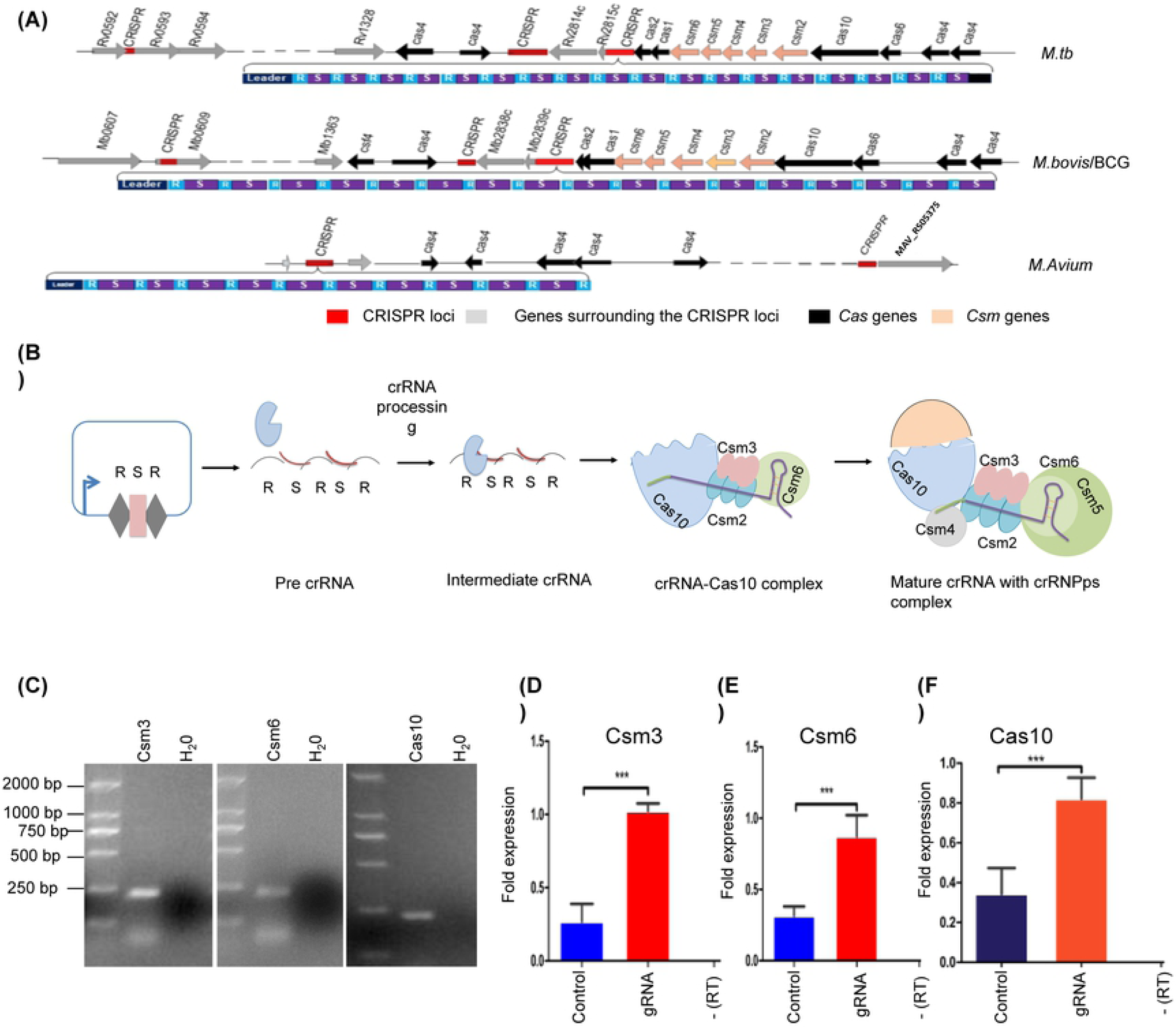
Schematic representation of the type III-A CRISPR-Cas10 system loci, crRNA biogenesis, and Cas gene expression in the *Mycobacterium tuberculosis* complex. **(A)** Type III-A CRISPR-Cas10 system in *M.tb, M.bovis*, and *M.avium*. **(B)** Schematic show of crRNA biogenesis by the type III-A CRISPR crRNA-csm complex. This crRNA-csm coplex then cleavage RNA or DNA through a transcription dependent manner **(C)** Expression of Csm3, Csm6 and Cas10 in *M.tb*. **(D-F)** Real-time PCR identification of the expression of Csm3, Csm6 and Cas10 in wild-type and gRNA-overexpressing strains.

To investigate the expression of type III-A CRISPR system genes in *M.tb*, we performed qRT-PCR with three functional candidate genes (*csm3, csm6*, and *cas10*). As shown in Figure 1.C, *csm3, csm6*, and cas10 are constitutively expressed in *M.tb* (primers are listed in Table S2). To determine whether the expression of the CRISPR array can stimulate the expression of this *cas* genes in *M.tb*, we constructed an expression plasmid (pMV-261 psmyc crRNA) harboring two 36-bp repeat sequences from CRISPR locus 3 were synthesized and cloned into pMV-261 to generate pMV-261-crRNA. Next, a 40-bp gRNA was cloned in between the spacers in the pMV-261-crRNA plasmid (Figure 1.B). Notably, the expression levels of these three genes increased significantly when *M.tb* were transformed with the plasmid expressing the mini-CRISPR array under the control of Psmyc promoter compared to wild-type strain (Figure 1.D-F). The Sanger sequencing of the PCR amplicon confirmed the sequence of the three genes. (Figure S2.E). A (-RT) control was also used to ensure that there is no genomic DNA contamination. Together, these results suggested that the expression of *cas* genes was enhanced upon the expression of CRISPR array.

As the type III-A CRISPR system can cut single stranded DNA (Kazlauskiene et al., 2016), we harnessed this system for *M.tb* self-DNA targeting by designing a 40-bp gRNA complementary to the coding strand of the target region and the corresponding homology directed repair (HDR) template (Figure S1. A and C, Table S1). The gRNA and HDR templates were cloned into the pMV-261-crRNA plasmid. The plasmids containing gRNA plus HDR, gRNA only or HDR only were then transformed into *M.tb*, respectively. A fourth plasmid without any gRNA or HDR (vector control) was also transformed into *M.tb* as a control. Figure S5.A demonstrate a significant reduction in cfus of the bacteria transferred with only gRNA plasmid compared to the bacteria transformed with gRNA plus HDR. Notably, there was still a significant reduction in the number of cfus of *M.tb* transformed with gRNA plus HDR as compared to the bacteria transformed with vector control (Figure S5.A). These data suggesteded that the self-DNA targeting caused a certain level of lethality to the bacteria but can be partially rescued by HDR template.

### Gene knock-in in *M.tb using* the type III-A CRISPR-Cas system

As *M.tb* contains an active endogenous type III-A CRISPR-Cas system, we next aimed to apply this system for gene knock-in in *M.tb* (Figure 2.A). First, we aimed to introduce a *GFP* gene into the *M.tb*H37Ra genome and construct a gyrA-GFP fusion protein strain using a 40-bp gRNA that was complementary to the coding strand of *gyrA* (Table S1). To facilitate site-specific recombination, an HDR template containing a *GFP* gene as a selection marker was inserted into the pMV-261-gRNA*-gyrA* and then transformed into *M.tb*. As shown in Figure 2.B, fluorescence was observed in colonies transformed with the plasmid containing the HDR template plus gRNA but not in colonies transformed with the HDR template only. Upon examining the fluorescence of multiple colonies (100), we observed fluorescence in more than 80% of colonies. To confirm insertion of the *GFP* gene in the correct position into the *M.tb* genome, the fluorescent colonies were boiled for PCR identification. The PCR was performed by using the primer pairs F/R and F/R1 (Figure S1.B, Table S2) yielded bands that were 1.72 and 1.1kb in size, respectively, indicating that the GFP had been knocked-in (FigureS3.A). This finding was further confirmed by Sanger sequencing of the PCR products (Figure 2.C). Together this data demonstrating that the type III-A CRISPR system can be used for gene knock-in in *M.tb*.

**Figure 2:**
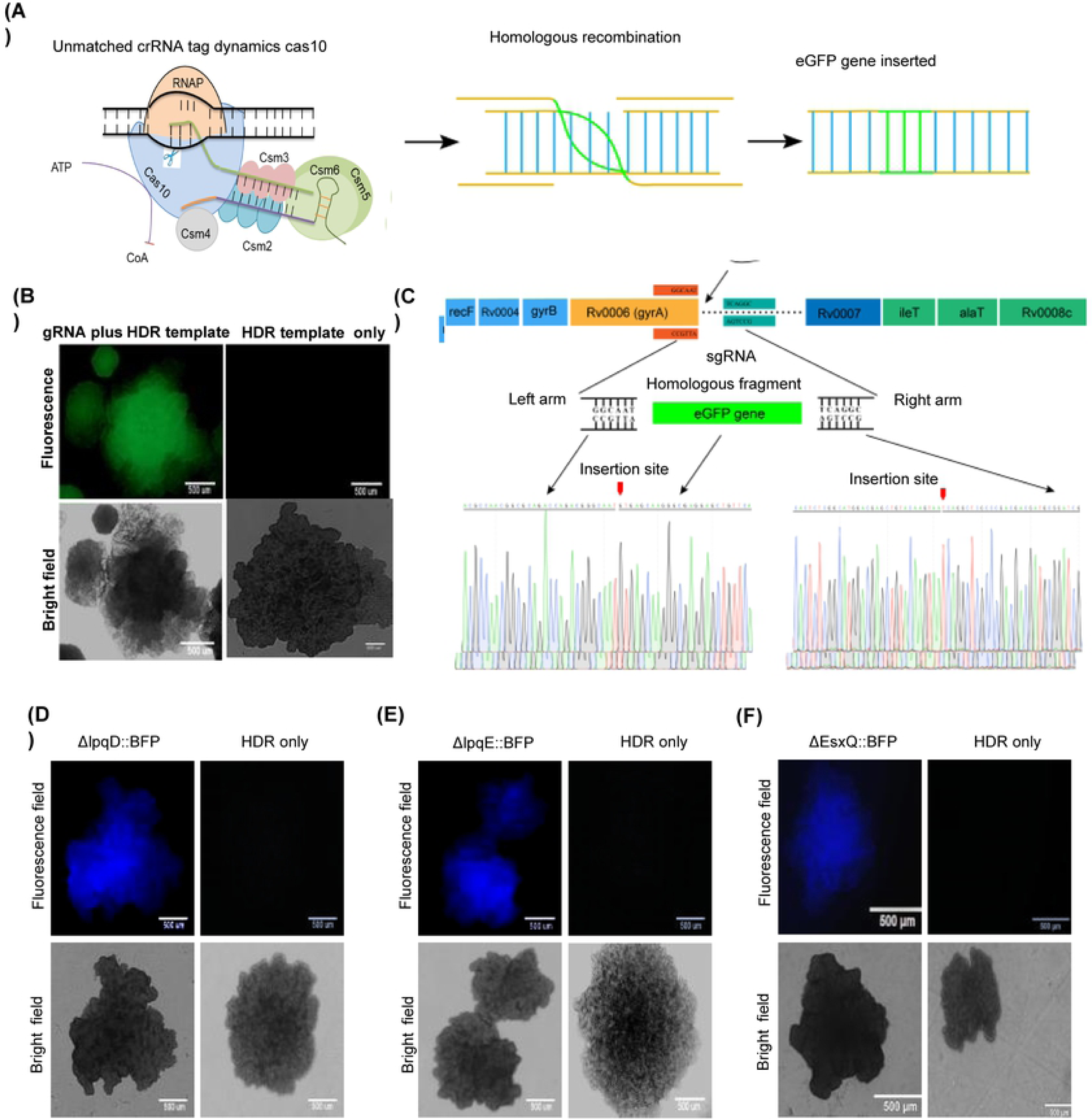
Endogenous CRISPR-Cas10 system-mediated gene knock-in and out in *M.tb*. **(A)** *M.tb* gene editing strategy in which CRISPR-Cas10-mediated specific DNA cleavage through a transcription dependent manner that can facilitates gene knock-in/out. **(B)** Green fluorescence in bacteria with the GFP gene was integrated after the *gyrA* gene locus by the endogenous CRISPR-Cas10 system. Fluorescence was lacking in the control group transformed with the HDR template only. **(C)** Cartoon representation of GFP insertion in the locus after *gyrA* and confirmation of insertion by Sanger sequencing. **(D-F)** Fluorescence and bright-field images of the colonies of *lpqE, lpqD*, and *esxQ* deleted strains.

### Gene knock-out using the type III-A CRISPR-Cas system

Next, we attempted to knock out 9 genes belonging to the early secretory family and lipoprotein family via the type III-A CRISPR system as a proof of concept. For this purpose a 40-bp gRNAs targeting the coding strands of the corresponding genes (Table S1) were cloned into the pMV-261-crRNA plasmid. As a selection marker, the *BFP* or *GFP* gene fused with the corresponding HDR template arms was cloned into plasmids containing gRNAs and then transformed into *M.tb*. Fluorescence was observed in colonies transformed with the plasmid containing the gRNA and HDR template arms but not in the control colonies (without gRNA) (Figure 2.D-F and Figure S3.A).To further confirm the knock-out of the target genes (*lpqN. lpqE, lpqD, esxA etc.*), the fluorescent colonies were selected for PCR amplification with the specific primers listed in Table S2.The PCR of Δ*lpqD*::BFP and *ΔesxQ*::BFP yielded a product with the expected size (1.2 kb), suggesting replacement of respective genes by *BFP* gene (Figure S3.B and D). A 1.75 kb band, indicating Δ*lpqE*::BFP, was amplified from the colonies expressing BFP, while amplification of the control (plasmid with only gRNA and or only HDR template) bacilli yielded a 1.5 kb band (Figure S3.C and E). The PCR products were further subjected to Sanger sequencing, which confirmed the replacement of target genes by the *BFP* gene (Figure S2.B-D).).

### Efficiency and off-target rate evaluation of endogenous type III-A CRISPR-Cas system-mediated gene editing

To evaluate the off-target effects of the type III-A CRISPR-Cas system, whole-genomic DNA libraries were constructed for the wild-type and H37Ra *ΔesxQ*::BFP, Δ*lpqD*::BFP, *ΔRv0839*::BFP strains and subjected to high-throughput sequencing (at least 250× for each sample) (Figure 3.A) The raw data were trimmed using Trimmomatic (Bolger et al., 2014), and the clean reads were mapped with the *M.tb*H37Ra reference genome using BWA (Li and Durbin, 2009) (Figure 4.B). The off-target rate was examined by SAMtools (Li et al., 2009) and BEF tools (Danecek and McCarthy, 2017). The *BFP* gene has been inserted at the target position (Figure 3.C, S4.C & D) without any potential off-target Indels in the genomic regions which includes gRNA and 20bp flanking region (80 bp in total) (Figure 3.D and E).

**Figure 3:**
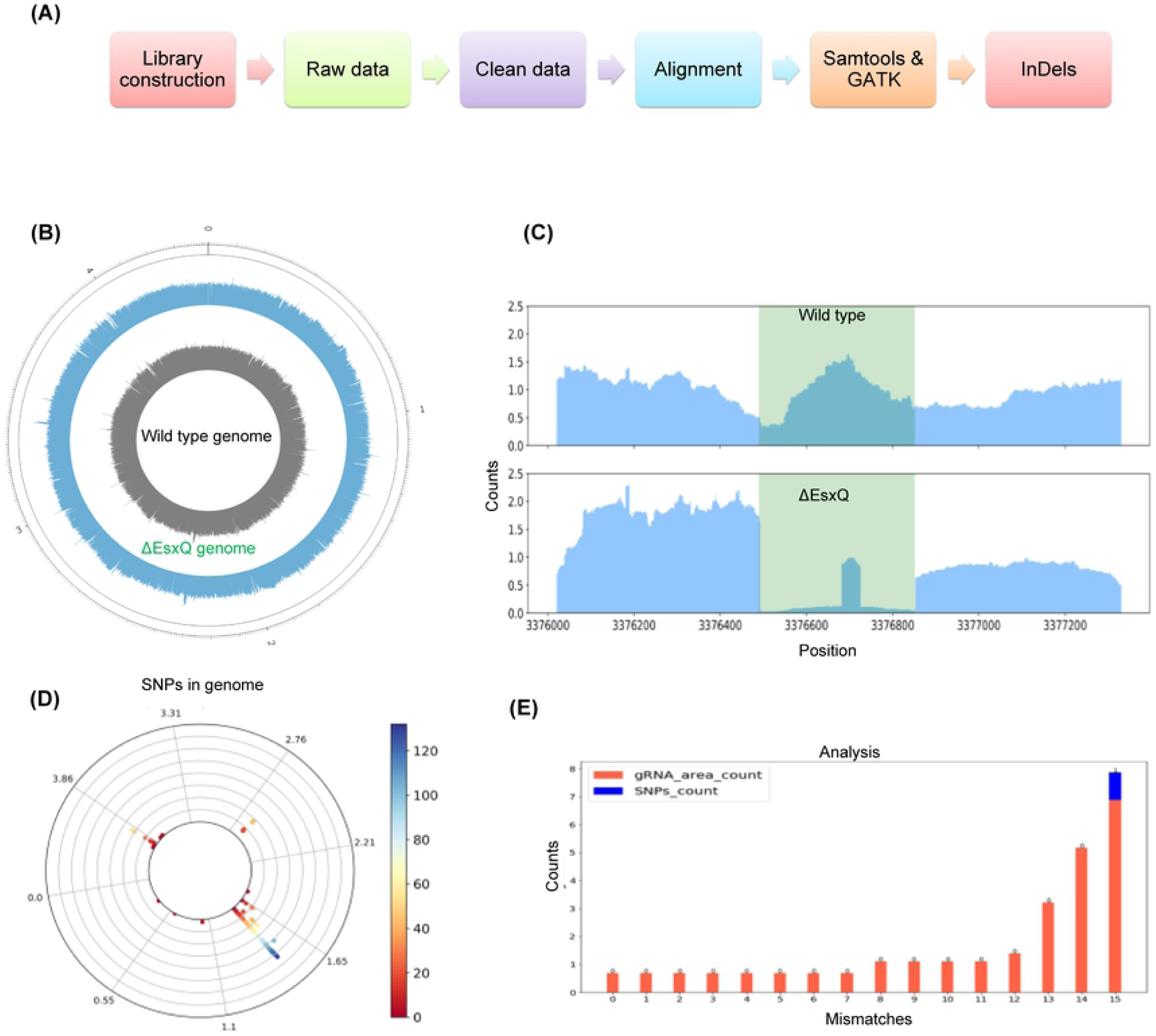
Gene knock-out and off-target effect analysis by whole-genome sequencing. **(A)** Schematic diagram for the gene knock-out and off-target effect analysis from DNA library preparation to in silico analysis. **(B)** Circos plot showing the genomic outline of the wild-type and esxQ knock-out mutant strains. **(C)** Representation of the sequencing reads aligned on the esxQ locus in the wild-type and ΔesxQ strains. **(D)** The polar chart represents genome-wide SNPs in the Δ*esxQ* strain compared to the wild-type strain. The colors indicate the number of SNP events at the specific sites, and the circle represents the whole genome. **(E)** Whole-genome off-target effect evaluation in the ΔesxQ strain with different mismatch cutoff values. No off-target effect was observed in the locus considered the gRNA length flanking by 20 bp region at both side.

**Figure 4:**
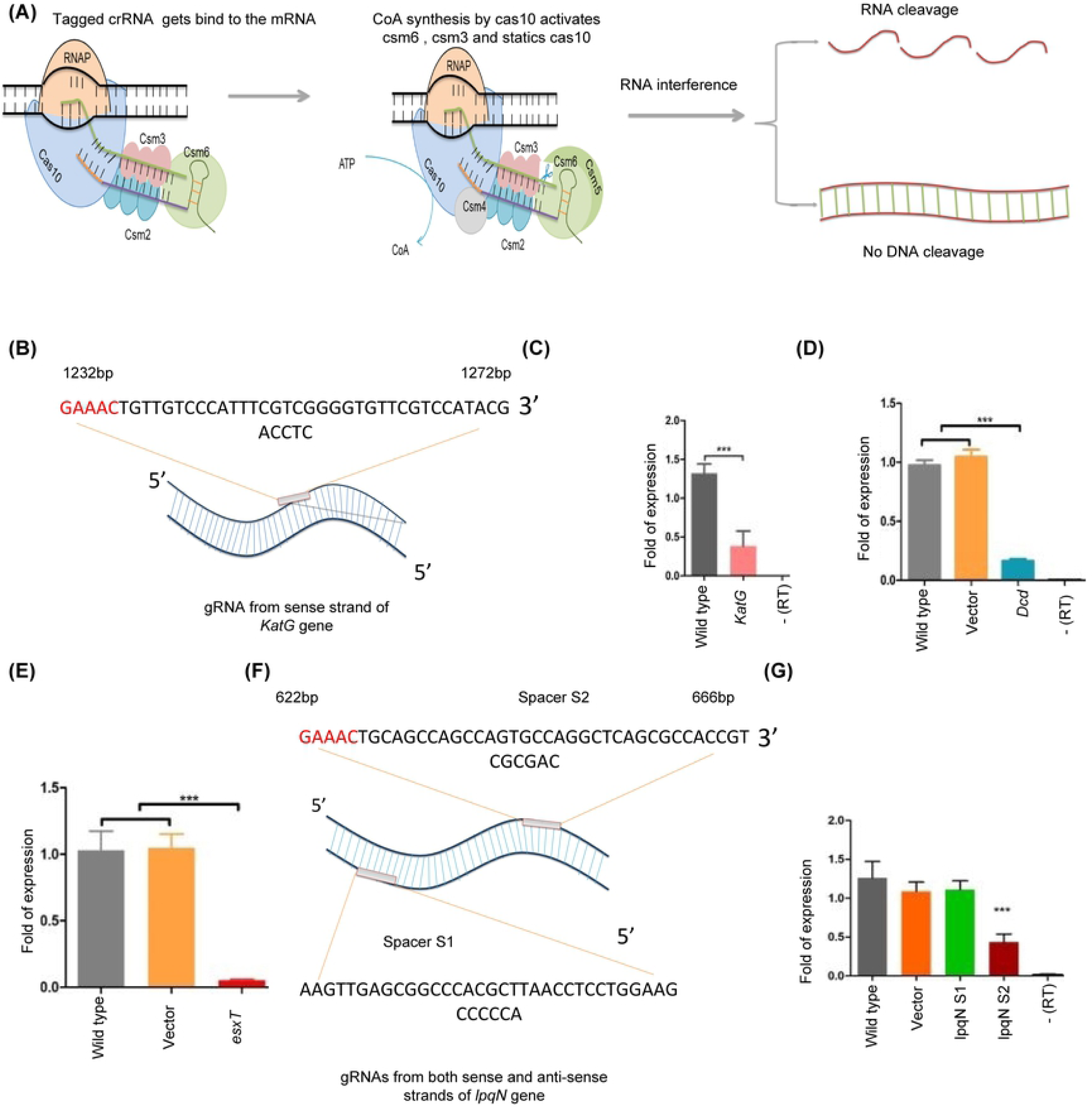
Endogenous CRISPR-Cas10-mediated gene interference in *M.tb*. **(A)** Schema for the CRISPR interference strategy in which the targeting sequence was flanked with a 5’-GAAAC-3’ tag, this tag recognize by the CRISPR cas complex which stop the nuclease activity of cas10 and switch it to synthesize coA (cyclic oligoadenylate) from ATP which leads to the activation of csm3 and csm6 resulting in RNA cleavage without DNA targeting. **(B)** Map and position of the gRNA for the *katG* gene. The pentanucleotide motif at 1227-1232 bp on the noncoding strand of *katG gene* along with the flanking region (40 bp) from 1232-1272 bp as potential gRNAs. **(C)** Inhibition of *katG* expression in the *katG* interference strain in comparison to the wild-type control strain. **(D and E)** Fold change in *dcD* and *esxT* gene expression in the *dcD* and *esxT* interference strains, respectively, in comparison to the vector control strain and the wild-type control strain. **(F)** Position of the gRNA on the coding and noncoding strands of the *lpqN* gene, designated as S1 and S2, respectively. **(G)** Inhibition of *lpqN* gene expression in comparison to the vector, S1 gRNA and wild-type control strains (*P < 0.01; ***P < 0.0001).

To check the genome editing efficiency of this system, *M.tb* H37Ra was transformed with plasmids containing the gRNA and HDR template corresponding to *Rv0847* (*lpqE*) and two control plasmids (one containing the HDR template only but no gRNA and the other containing the gRNA without the HDR template). Fluorescence was observed in approximately 80% of the colonies transformed with the plasmids containing the gRNA plus HDR template, while no fluorescence was observed in the control (Figure 2E). To further verify the efficiency, 11 randomly picked colonies were cultured for 5 to 7 days and amplified with the primers of *lpqE* KO detection F and R (see Table S2 for primer information). As shown in Figure S3.F, 10 clones out of 11 exhibited insertion of BFP, while 1/11 sample showed a same band with wild type strain. The Sanger sequencing of these PCR bands showed that the *lpqE* gene was replaced by BFP (Figure S2.D).

### Harnessing the endogenous Type III-A CRISPR system for CRISPR interference in *M.tb*

Next, we aimed to reprogram type III-A CRISPR system for gene interference in *M.tb.* To this end, a 40 bp gRNA on the noncoding strand flanked by the pentanucleotide 5’-GAAAC-3’, matching the repeat handle sequence, was selected as a target sequence (Figure 4.A). This sequence would mimic the endogenous CRISPR array and thus circumvent DNA cleavage but allow RNA interference. Accordingly, a 40 bp gRNA in the *katG* gene flanked by the pentanucleotide motif at 1228-1232 bp (5’-GAAAC-3’) was used as a gRNA to specifically target the RNA of the *katG* gene without any DNA interference(Figure 4.B). The gRNA (Table S1) was cloned into pMV-261-crRNA and transformed into *M.tb*. The qPCR results showed that the expression of *katG* was significantly inhibited compared to that of the wild-type control (Figure 4.C). Two genes, namely, *dcd* (dCTP deaminase) and *esxT*, were used to further investigate the CRISPRi of the gene using the same strategy. The 40 bp gRNA flanked by 5’-GAAAC-3’ in both genes was selected (TableS1) and cloned into pMV-261-crRNA. pMV-261-crRNA without a gRNA was also included as a plasmid control. The qRT-PCR results showed that *esxT* and *dcd* significant different compared to the wild type and plasmid control (Figure 4.D and E).

To determine whether RNA interference could be achieved by targeting the noncoding strand, we selected the *lpqN* gene as a target. Two 40-bp gRNAs, designated spacer-S1 (targeting the noncoding strand) and spacer-S2 (targeting the coding strand), of the *lpqN* gene were then inserted into pMV-261-crRNA and then transformed into *M.tb* H37Ra (Figure 4.F). The gRNA targeting the noncoding strand (spacer-S1) should not bind to the mRNA and thus should theoretically be unable to inhibit the expression of the target gene. Consistent with this hypothesis, the qRT-PCR results showed that the mature gRNA targeting the *lpqN*-coding strand (spacer-S2) efficiently inhibited the expression of the gene, whereas no significant downregulation was observed for spacer-S1, the plasmid and the wild-type control (Figure 4.G). Taken together, these results suggest that the type III-A CRISPR-Cas system can efficiently inhibit gene expression by targeting the coding strand.

To confirm that the system is successfully binding to and cuts the target mRNA, we performed RNA immune precipitation qPCR (RIP-qPCR) assay. The CRISPR complex was purified through csm6-HA tag from the bacteria containing RNA targeting plasmid for *kaG* and *lpqE* genes respectively (Figure S5.G). Upon normalization to negative control, we found that the groups with csm6-HAtag plus gRNA were highly enriched in *katG* and l*pqE* mRNA as compare to the groups containing no gRNA (Figure S5.E). To confirm the RNA interference is not due to the cutting of the template DNA, we amplified *katG* gene from *katG* RNA interference strain for Sanger sequencing, which demonstrated no mutation in *katG* gene (Figure S2.F). To validate this phenomenon, two more different gRNAs targeting *katG* DNA and RNA, were cloned into pMV-261-crRNA plasmid, respectively (table S1) and transformed into *M.tb* individually. A pMV-261-crRNA plasmid without any gRNA was also transformed into *M.tb* as a control. Figure S5.F showed that the number of cfus was highly reduced in the DNA targeting group as compared to the RNA targeting group or empty plasmid group. Of note, there was no significant difference between the cfus of RNA targeting group and empty plasmid group. These data demonstrated that the gRNAs designed for RNA interference may lack the DNA cutting. To check the polar effects of type III A CRISPR system mediated RNA interference, we performed qRT-PCR for the neighbor genes of *lpqE* and *dcD* knocked down genes respectively. Figure S5.B & C showed little polar effect on the neighbor genes in case of *lpqE* knocked down, while there was a slight polar effect of *dcD* knocked down gene..

### Simultaneous multiple-gene interference in *M.tb* by the endogenous type III-A CRISPR

In the endogenous type III-A CRISPR-Cas system, the CRISPR array comprising multiple spacers flanked by repeats is transcribed into a long primary transcript known as the precursor-crRNA (pre-crRNA). This RNA is processed into its mature form by the endogenous Cas enzyme, yielding multiple mature crRNAs that bind to the corresponding sites on mRNA, causing cleavage at multiple sites on targeted genes (Figure 5.A). Therefore, we aimed to accomplish simultaneous silencing of multiple genes by a single plasmid using type III-A CRISPR-Cas as an interference system. Three spacers targeting the *lpqE, katG* and *inhA* genes flanked by repeats on both sides were then synthesized and cloned into pMV-261-crRNA and transformed into *M.tb* (Figure 5.B). The qRT-PCR results demonstrated that the expression of these three genes was simultaneously downregulated with a high efficiency compared to that of the control (Figure 5.C), indicating that the endogenous type III-A CRISPR-Cas system can be employed for simultaneous silencing of multiple genes with high efficiency.

**Figure 5:**
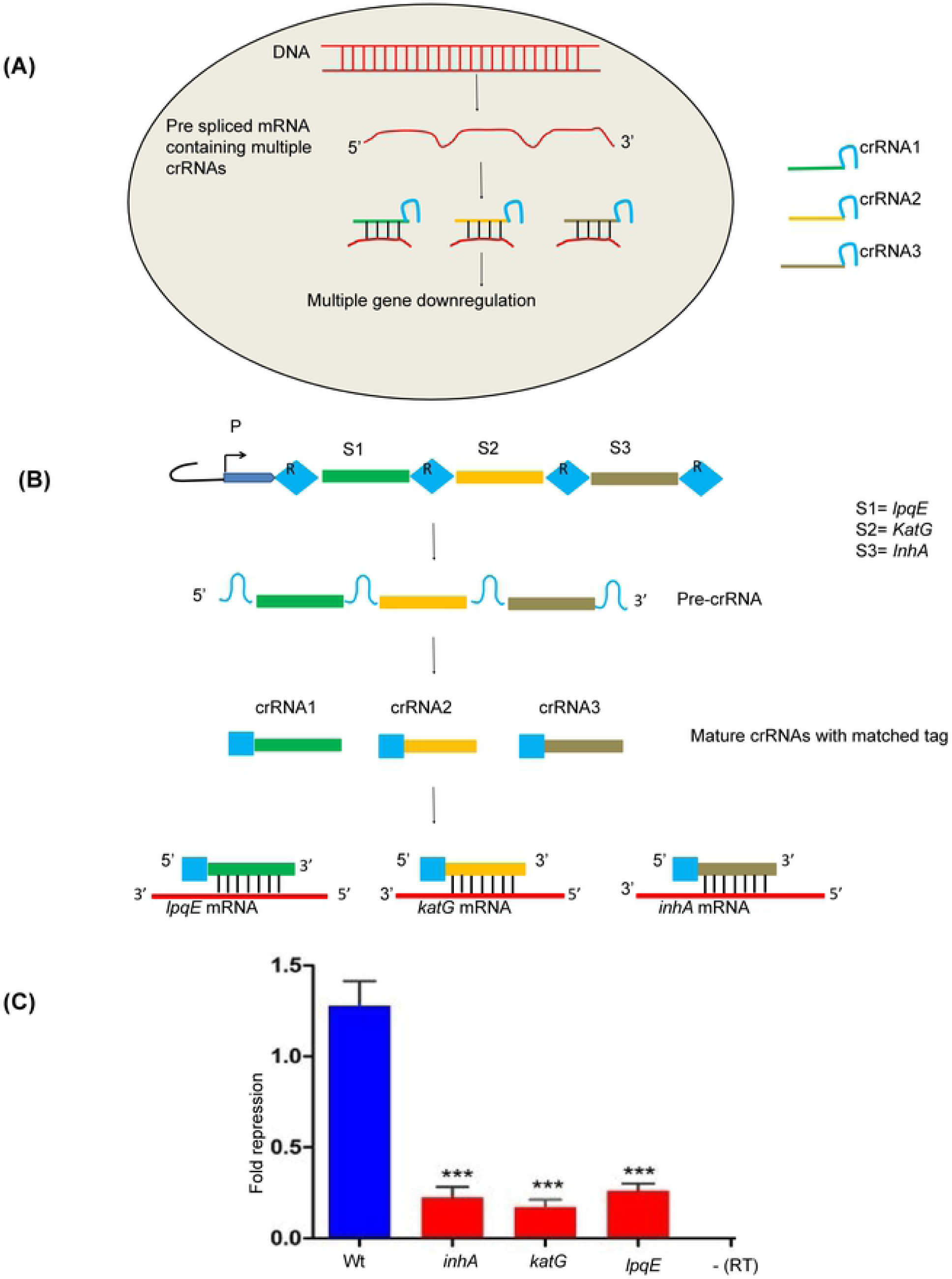
Schema for endogenous CRISPR-Cas10 mediated simultaneous multiple-gene interference in *M.tb*. **(A)** Binding of multiple crRNAs to the corresponding targeting mRNAs. **(B)** Schema for the expression of multiple spacers, generation of multiple crRNAs and the binding of the mRNAs to the corresponding target mRNAs. **(C)** Simultaneous inhibition of *lpqE, katG* and *inhA* genes expression by the three spacers in comparison to the wild-type strain. Significant difference is indicated by the asterisks above the bars (***P< 0.0001).

### Genome-wide CRISPRi screening for *M.tb in-vitro* growth-regulating genes

Next, we aimed to accomplish genome-wide CRIPSRi screening based on the reprogrammed type III-A CRISPR-Cas system. For this purpose, a library pool containing 5658 gRNA fragments (Table S3) was generated via on-chip oligo synthesis and cloned into the pMV-261-crRNA plasmid. This library was first transformed into *E.coli* for plasmid amplification and gRNA input composition analysis. After amplification, this gRNA library was electroporated into *M.tb* for screening growth-regulating genes (Figure 6.A).

**Figure 6:**
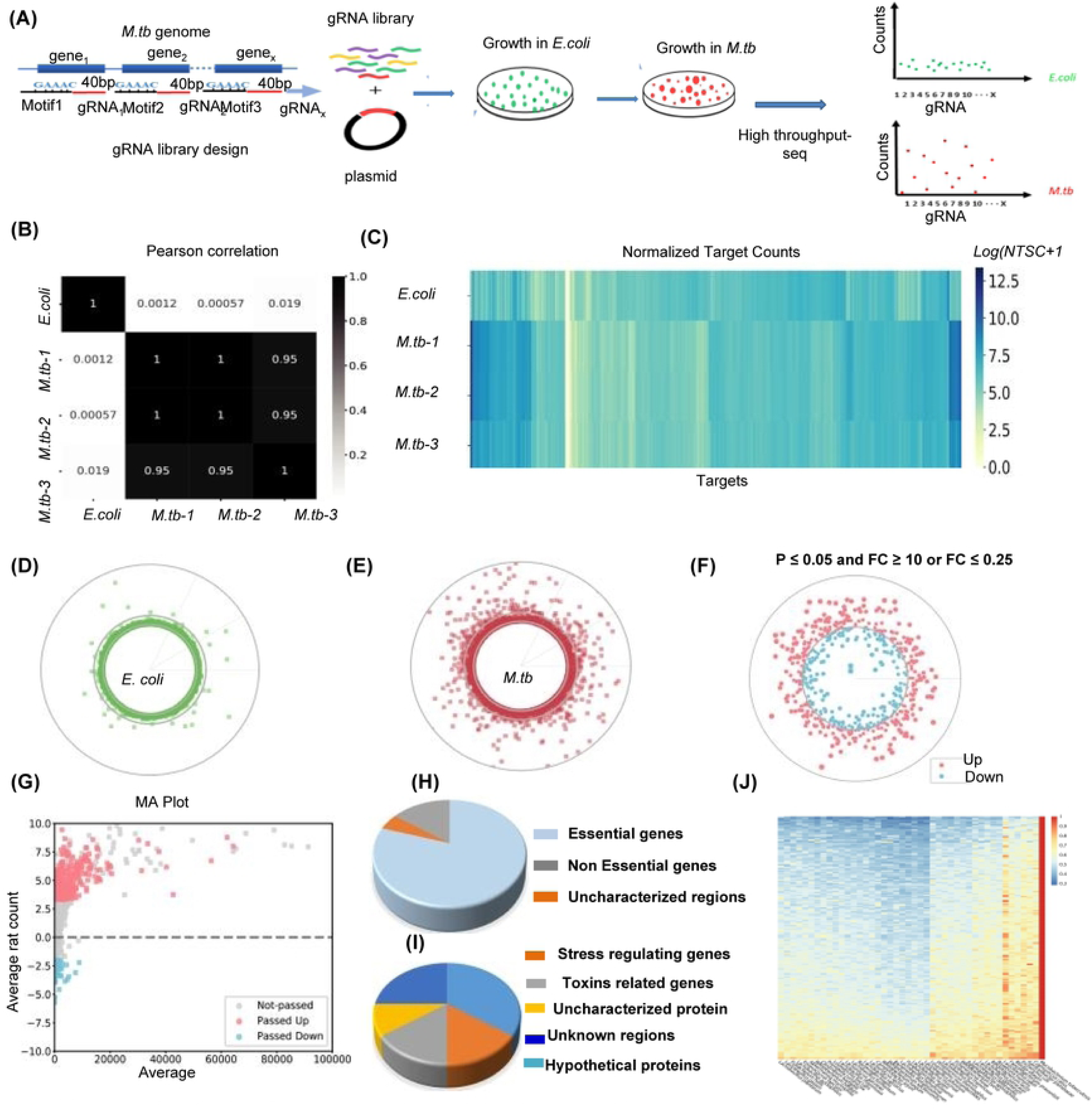
Genome-wide CRISPR interference screening for growth-facilitating genes in *M.tb*. **(A)** Schematic representation gRNA library construction and screening: 5658 gRNAs were synthesized and cloned into *E.coli* and then subjected to high-throughput sequencing, for which the gRNA read count distribution was used as input. The gRNA library was electroporated into *M.tb* to interfere with target gene expression. gRNAs targeting growth-regulating genes influence *M.tb* growth, which can be revealed by high-throughput sequencing and analysis of the gRNA read counts from the *M.tb* plasmids in comparison with that of the input from *E.coli.* **(B)** Pearson correlation of gRNA sequence count distribution of three *M.tb* samples with *E.coli* (input). **(C)** Heat map of the gRNA read count of three *M.tb* samples in comparison with the *E.coli* input. **(D-E)** Global distribution of gRNA reads in the *E.coli* genome and *M.tb* genome (mean values). Each point represents a gRNA, and the radius represents the read count. **(F)** Circle plot of the CRIPSRi screening result. The red points and blue points represent gRNAs with significantly increased and decreased levels (*M.tb* vs *E.coli*), respectively, aligned to the corresponding target genes in the genome. The distance to the inner circle of each point represents the fold change value. **(G)** MA plot of the CRIPSRi screening result shown in (F). The x-axis shows the average gRNA read count, and the y-axis shows the log2 fold change (*M.tb* vs *E.coli*) values. (H) Pie chart showing the properties of the top 50 low-counted target genes, among which 78% have been previously reported to be essential for growth. **(I)** Percent of different types of genes among the top 50 high counted gRNAs. **(J)** Heat map representing the similarity of the 208 growth-facilitating genes between *M.tb* and the 43 most common probiotics.

As a proof of concept, we collected approximately 1.5 million *E.coli* colonies and prepared a mixed gRNA library construct. This library was aliquoted and electrotransformed into three independent competent *M.tb*. After 3 to 4 weeks of incubation, a total of approximately 1.5 million *M.tb* CFUs from 330 plates were harvested for plasmid preparation. The gRNA were amplified from *E.coli* and *M.tb* plasmids by PCR and subjected to library construction for high-throughput sequencing.

We hypothesized that the *M.tb* gRNAs with significantly reduced counts compared to *E.coli* gRNAs would likely target growth-facilitating genes, while the gRNAs with significantly increased counts probably targeted growth-repressing genes. We first screened the reads matched with repeat-gRNA-repeat structure and then trimmed the target sequences for further analysis. The counts of each target sequence were normalized using the median-of-ratios method (Rousset et al., 2018). The high Pearson correlation coefficient between the three independent *M.tb* gRNA libraries indicated that the composition and distribution of the target sequences and the corresponding read counts were highly similar; therefore, the screening assay was highly reproducible (Figure 6.B-C). Next, we analyzed the normalized count distribution of each target in the *E.coli* and *M.tb* libraries. The count distribution of most of the targets in *E. coli* library was equally distributed, whereas the counts of many targets in the *M.tb* library were highly scattered (Figure 6.C-E), suggesting that many of the gRNAs influenced *M.tb* growth.

We further screened the gRNAs with significantly reduced or increased count numbers in the *M.tb* library compared to the *E.coli* library (Figure 6.F and G, Table S4). Finally, we identified 228 genes as potential *M.tb* growth-facilitating genes and 385 genes as potential *M.tb* growth-repressing genes (Figure 6.F and G, table S4.H). Notably, among the top 50 potential growth-facilitating genes, 39 (78%) have been reported to be essential for growth (DeJesus et al., 2017; Griffin et al., 2011; Sassetti et al., 2003; Zhang et al., 2012), which supports the reliability of this CRISPRi screening method (Figure 6.H). Among the top 50 potential growth-repressing genes, we found that 15% belonged to the toxins family, 35% were stress-related proteins, 10% contained unknown regions, 15% were hypothetical proteins, and 25% were uncharacterized proteins (Figure 6.I).

Because genes that facilitate or are essential for growth could potentially be used as anti-*M.tb* drug targets, we next analyzed the similarity between the 208 growth-facilitating genes from *M.tb* and the whole proteomes of 43 probiotics. The top 31 genes in the heat map in Figure 6.J exhibited the least similarity with the whole proteomes of 43 probiotics. This data suggest that these genes could be used as a drug target with conceivable fewer side effects on probiotics than other genes and could be ideal anti-*M.tb* drug targets (Table S5).

### Genome-wide CRISPRi screening for *M.tb* intracellular growth-regulating genes

Finally, we aimed to apply this system to perform genome-wide CRIPSRi screening for growth-regulating genes during intracellular growth (growth inside macrophages). To this end, a total of approximately 1.5 million *M.tb* CFUs containing 5658 gRNA library in triplicate were infected with Thp-1 cells and grew for 3 days after removing the extracellular bacteria. As a reference, the *M.tb* containing 5658 gRNA library in triplicate were cultured *in-vitro* in Middlebrook 7H9 broth (Figure7.A). In this assay, the *M.tb* gRNAs with significantly reduced counts during intracellular growth would likely to be the genes that are highly required for intracellular *M.tb* growth (Figure 7.A).

**Figure 7:**
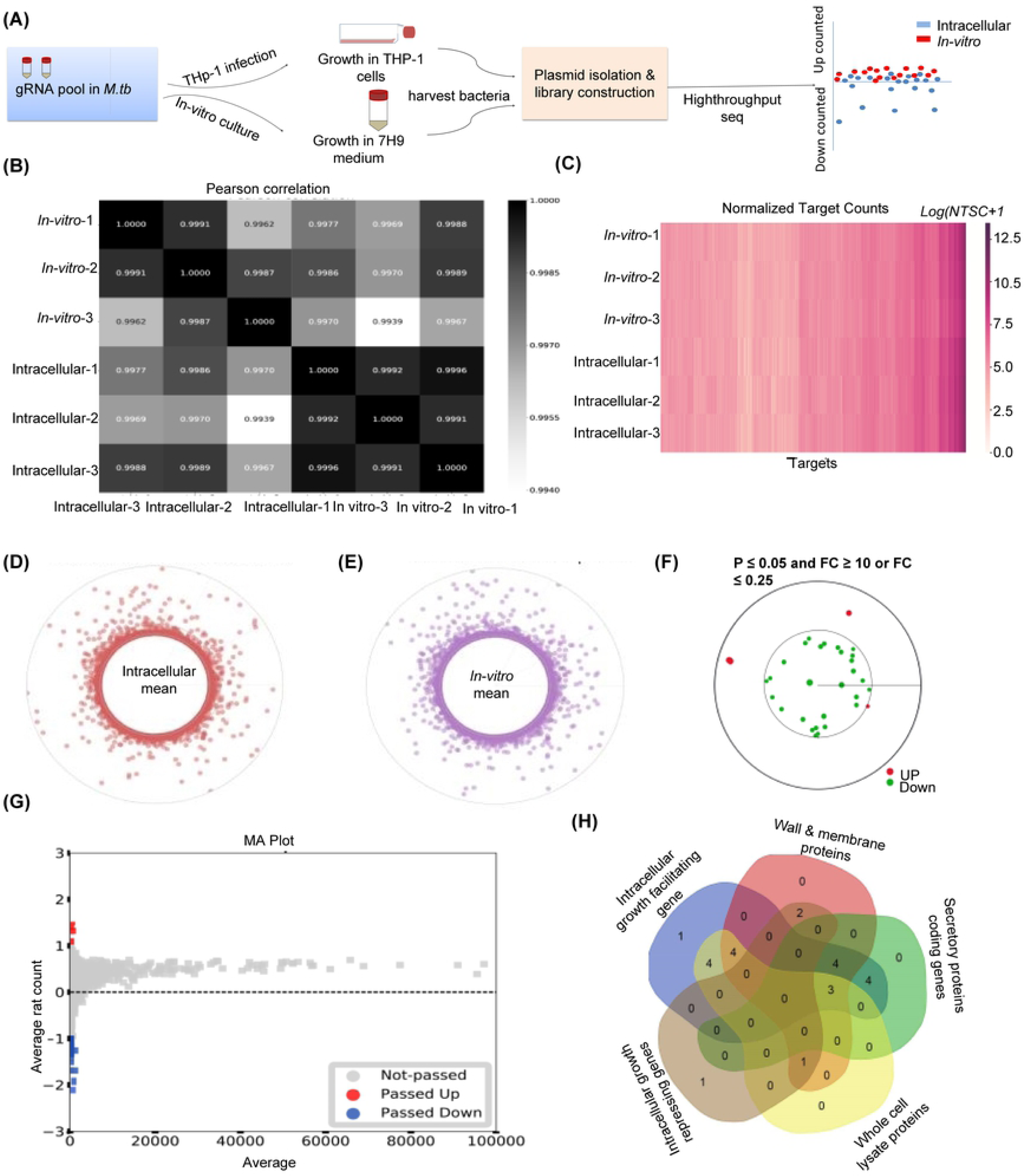
Genome-wide CRISPR interference screening for *M.tb* intracellular growth-regulating genes. **(A)** Schema for Genome-wide CRISPR interference screening for *M.tb* intracellular growth-regulating genes. **(B)** Pearson correlation of gRNA sequence count distribution of three independent *in-vitro M.tb* cultures with intracellular *M.tb*. **(C)** Heat map of the gRNA read count of three independent *in-vitro* cultures in comparison with the intracellular *M.tb*. **(D-E)** Global distribution of gRNA reads in the *in-vitro* and intracellular *M.tb* (mean values). Each point represents a gRNA, and the radius represents the read count. **(F)** Circle plot of the CRIPSRi screening result. The red points and blue points represent gRNAs with significantly increased and decreased levels (*in vitro* vs inside THp-1), respectively, aligned to the corresponding target genes in the genome. The distance to the inner circle of each point represents the fold change value. **(G)** MA plot of the CRIPSRi screening result shown in (F). The x-axis shows the average gRNA read count, and the y-axis shows the log2 fold change (*in vitro* vs inside THp-1) values. **(H)** Venn diagram represents different category of the intracellular growth-regulating genes.

The gRNA from both *in vitro* culture and intracellular bacteria was amplified and subjected to library construction for high-throughput sequencing. The raw reads were screened according to the repeat-gRNA-repeat structure and then normalized using the median-of-ratios method (Rousset et al., 2018). As shown in Figure 7.B, the high Pearson correlation coefficient between the three independent *M.tb* gRNA libraries indicated that the composition and distribution of the target sequences and the corresponding read counts were highly reproducible. Next, we analyzed the normalized count distribution of each gRNA target in the *in-vitro* and intracellular groups (Figure 7.C-E). Furthermore, the gRNAs with significantly reduced or increased numbers in the intracellular groups compared to the *in-vitro* group were screened (Figure 7.F and G). Finally, we identified 29 highly lower count genes as potential *M.tb* intracellular growth-facilitating genes and 4 highly counted genes as potential intracellular growth-repressing genes (Table S8). Among the top 20 highly growth essential genes, 5 genes were coding for secretory proteins, 11 genes belong to the membrane and cell wall component proteins while 4 belong to the cytosol and membrane proteins (de Souza et al., 2011; Mawuenyega et al., 2005; Xiong et al., 2005) (Figure 7.H)

## Discussion

In this study, we reprogrammed the endogenous type III-A CRISPR-Cas system to develop a robust and versatile tool for gene knock-in, gene knock-out, gene interference and CRIPSRi screening in *M.tb* by simple delivery of a plasmid containing the mini-CRISPR array. We demonstrated that the type III-A CRISPR-Cas system can dramatically enhance the site-specific recombination frequency, the positive rate of which can reach up to 80%. In addition to high efficiency, this system has the advantage of a simple procedure, requiring only two-step cloning (first of the gRNA and then of the cognate donor DNA). In this regard, the currently, used gene editing tool in *M.tb-*based phage transduction is laborious and time consuming due to the multiple cloning steps involved and the screening for positive phages; in addition this tool requires expensive kits and expertise (Bardarov et al., 2002; Choudhary et al., 2016).

The type III-A and B systems exert an anti-plasmid immunity effect by degradation of the plasmid DNA and the target transcript (Deng et al., 2013; Samai et al., 2015). The type III-B system has been used for gene editing and gene silencing in *Sulfolobus islandicus* (Li et al., 2015; Peng et al., 2015). Recently, Guan et al. reported the first application of the type III-A system for gene deletion (Guan et al., 2017).To the best of our knowledge, the present study is the first to demonstrate the type III-A CRISPR-Cas system-mediated “killing two birds with one stone” tool for robust gene editing and silencing in *M.tb*.

Genetic manipulation, especially of essential genes in *M.tb*, is crucial for understanding gene function and identifying targets for anti-*M.tb* drugs and vaccines. A codon-optimized gene interference approach based on the dCas9 system has been developed to silence specific genes in *M.tb*, which can reduce gene expression by approximately 4-fold (Choudhary et al., 2015; Singh et al., 2016). Recently, dCas9 from *S. thermophilus* was used for gene silencing in *M.tb* with reduced toxicity (Rock et al., 2017). However, these systems need to express the exogenous Cas9 proteins, making it difficult to avoid proteotoxicity. The endogenous type III-A CRISPR-Cas-based specific gene silencing system was obtained by simply transforming a plasmid expressing a mini-CRISPR array with a gRNA into *M.tb*.

After transcription, the mini-CRISPR RNA is reprogrammed by the endogenous Csm complex to bind with the mRNA of the target gene, thus inhibiting expression of the target gene. This system offers several advantages over the CRISPR-Cas9 system, such as the simplicity of cloning of the mini-CRISPR array and gRNA without introducing any exogenous *Cas9* gene, minimizing the toxicity of the system. Unlike Cas9, there is no stringent requirement of the PAM sequence for gene targeting. Similar to the endogenous type III-A CRISPR-Cas system, the reprogrammed CRISPR array can also simultaneously generate multiple spacers. These spacers can be processed into multiple mature crRNAs that bind to the corresponding target sites and inhibit the expression of multiple genes simultaneously. This approach can solve the problem of functional redundancy in functional genomic research on *M.tb. M.tb* causes the leading infectious disease in the world, the pathophysiology and functional genomics of this organism remain poorly understood, which greatly hampers the development of new anti-*M.tb* vaccines and drugs. Whole-genome screening via gene silencing based on the CRISPR-Cas system is a powerful approach and has been widely used to explore gene function (Rousset et al., 2018; Wang et al., 2018). Recently, Wet et al. applied CRISPR-dCas9 screens for functional characterization of transcription factors in *M.smegmatis* (de Wet et al., 2018). However, no study has been performed on the genome scale in *M.tb*. In this study, we identified 5656 and 5658 potential targets for the H37Rv and H37Ra strains, respectively, covering almost all the coding sequences of the *M.tb* genome. We used this gRNA library for whole-genome-scale screening of growth-regulating genes as a proof of concept. Using this approach, we identified 208 genes that facilitated growth and 385 growth-repressing genes from H37Ra.

Notably, among the top 50 identified growth-facilitating genes, 78% have been previously reported to be essential for growth, which supports the reliability of this screening system. As most of the CRISPRi-identified growth-facilitating genes are also essential for bacterial growth, these genes could be used as potential anti-*M.tb* drug targets. In this study, we further analyzed the similarity between these growth-facilitating genes of *M.tb* and the whole proteomes of 43 probiotics and identified 31 genes as potential anti-*M.tb* drug targets that possibly have fewer side effects on probiotics than the other identified genes. The identification of new drug targets could be of great importance for protein-structure-based drug design and screening, considering the current severity of antibiotic resistance in *M.tb*.

The CRIPSRi screening also identified a certain number of the toxin family genes and stress-related genes as potential growth-repressing genes. Interference of the toxins family genes could conceivably inhibit the growth of *M.tb* via attenuated repression of toxin proteins. Therefore, our database of growth-repressing genes may be used to screen for toxin-antitoxin family genes. In this regard, previously identified antitoxin genes were present in our database of growth-repressing genes, such as *dinX* and *vapB5* (Gupta, 2009). Furthermore, the knock-down of some stress regulons and other transcriptional factor regulators for non-replicating persistence, such as *devR/dosR* and *sigA*, could also increase the growth of *M.tb* (Gengenbacher and Kaufmann, 2012). Thus, our CRIPSRi screening method may contribute to the identification of genes involved in toxin-antitoxin regulation, stress regulation and transition from active *M.tb* to the persistent form. The procedure for this screening system requires only one step of gRNA library cloning and electrotransformation into *M.tb*, which is much simpler and cost effective than the procedures of other screening systems, such as transposon-based mutagenesis. Moreover, in contrast to gene knock-out, this CRISPRi system only knocked down gene expression; thus, this system can also be used for genetic screening and investigation of the functions of genes essential for growth.

In our study, we also applied this CRIPSRi system to screen the genes regulating growth in the host cells, by which we identified 29 genes as potential *M.tb* growth facilitating genes and 4 genes as potential *M.tb* intracellular growth repressing genes. Among the top 20 highly down regulated genes, 18 genes have been reported to be essential for growth inside macrophages (Akhter et al., 2008; Ma et al., 2018; McAdam et al., 2002; Monahan et al., 2001; Sassetti and Rubin, 2003; Venugopal et al., 2011; Ward et al., 2010), whereas 2 genes were novel *M.tb* intracellular (inside Thp-1 derived macrophages) growth-facilitating genes. As examples among the highly down regulated genes, the *Senx3* belongs to the two component regulatory system and *groEL1* gene that belongs to the membrane protein genes of *M.tb*. It has been demonstrated that both of these genes are required for survival of *M.tb* inside the macrophages but not *in-vitro* (Lamichhane et al., 2003; Ojha et al., 2005). As these genes are dispensable genes for the growth of *M.tb* during *in-vitro* growth but, highly required during intracellular survival, they could be the potential vaccine candidates for tuberculosis. In our study, we also identified intracellular growth repressing genes, such as *ponA-*1 gene, which is involved in stress regulation (Saxena et al., 2008; Talaat et al., 2004). It would be of great interest to further investigate whether the knock-down of this gene could prevent *M.tb* from going to the latency stage inside macrophages. The limitation of this endogenous CRISPR-Cas system-based gene silencing is the dependency of this method on the pent nucleotide motif for gene silencing, although the system can cover a majority of the *M.tb* genome. Further research is needed to determine whether a motif with less than 5 bp (i.e., a di-, tri- or tetra nucleotide motif) could also be efficiently used for gene silencing and DNA targeting. Moreover, understanding the structure of the Csm-crRNA complex, target binding and molecular mechanism of DNAs/RNAs involved in natural immunity for genetic manipulation of the type III-A CRISPR-Cas system could make this tool broadly applicable in related research fields.

In summary, we reprogrammed the *M.tb* endogenous type III-A CRISPR-Cas10 system for simple and efficient genome editing, gene interference and CRISPRi screening. This system greatly facilitates genetic manipulation with high specificity and efficiency and provides an opportunity for genome-scale gene interference-based screening in *M.tb.* This system can be extensively used to explore the functional genomics of *M.tb* and will contribute to basic research on *M.tb* biology and to the development of anti-*M.tb* vaccines and drugs.

## Acknowledgments

This work was supported by The National Key Research and Development Program of China (Grant No.2017YFD0500303), the National Natural Science Foundation of China (Grant No. C180501, 31602061), the Huazhong Agricultural University Scientific &Technological Self-innovation Foundation (Program No. 2662017PY105 2662017PY105), Doctoral Fund of Ministry of Education of China (Grant No. 131012). We’d like to thank Wuhan GeneCreate Biological Engineering Co., Ltd. for their technique support.

## Author Contributions

G.C and M.J conceived the idea. M.J and K.R performed experiments. W.X and W.L performed the bioinformatics analysis. X.C, L.D provided some experiment materials. M.J, K.R, G.C wrote the manuscript, with inputs from all other authors. Z.F.F, X.C, Z.H, M.A.N, N.P and R.T reviewed and edited the manuscript. All the authors discussed the results and commented on the manuscript.

## Legends

**Supplementary Figure 1:** (A) Vector map of the construct for CRISPR-mediated gene editing and interference. (B) Map of primers used to identify GFP knock-in, in which the forward and reverse primers are located 100 bp upstream and downstream of the left and right arms, respectively. (C) Sequences of different parts of the plasmid, such as psmyc, which represents the promoter required for expression of the crRNA; the gRNA sequence was cloned into the plasmid pMV-261 at the BbsI site between the two repeat; and T1, which represent the terminator. (D) Sequences of the spacer used for simultaneous multiple-gene interference.

**Supplementary Figure 2:** Supplementary Figure 2: Type III-A CRISPR-Cas system-based gene deletion in *M.tb*. (A) Fluorescent colonies expressing BFP as a selection marker representing gene knock-out after replacement of the target genes indicated in each panel. (B, C and D) Representative Sanger sequencing chromatographs revealing BFP and GFP insertion into the target sites. (E) Sanger sequencing chromatographs of qRT-PCR products of the respective cas*/csm* genes. (F) Sanger sequencing chromatographs of *katG* gene from the wild type and *katG* knocked down strain.

**Supplementary Figure 3:** PCR confirmation of gene knocked Out/In in *M.tb* mediated by the endogenous CRISPR-Cas10 system. (A) PCR Confirmation of EGF knock-In into the *gyrA* locu. (B-E) PCR confirmation of gene knock-Out. (F) DNA targeting efficiency of the type III-A CRISPR system in *M.tb*. PCR amplification showing successful insertion of BFP in 10 out of 11 colonies. The bands on the gel from 1 to 11 represent H37Ra transformed with the plasmid containing the gRNA and HDR template for lpqE; 13 and 14 are the clones transformed with the plasmid containing only the gRNA or HDR template individually. 15 is the WT control and 16 is the water control.

**Supplementary Figure 4** (A and B) Circos plots showing the genomic outlines of the *esxC* and *lpqD* mutants strains along with their wild type strains. (C and C) Representation of the sequencing reads aligned on the *esxC* and *lpqD* locus in the wild type and the respective mutant strains

**Supplementary Figure 5**: (A) Targeting of Self DNA by the endogenous type III-A CRISPR system directed by a gRNA designed from the *gyrA* gene. (B and C) polar effects of the endogenous type III-A CRISPR mediated RNA interference on the neighbor genes of *lpqE* and *dcD* genes. (D) RNA immune precipitation assay performed for the detection Type III-A CRISPR complex binding to the target mRNA. (E) qRT-PCR based enrichment of the *inhA* and *lpqE* mRNA from the immune precipitated RNA. The fold enrichment has been normalized to the samples with no gRNA and no tagged csm6 samples. Then the normalized values of each sample were compared to the respective controls. (F) Effect of the type III-A CRISPR assisted RNA and DNA targeting on the cfus of *M.tb.*

**Supplementary Figure 6**: Strategy for gRNA library design for genome wide CRISPRi Screening in *M.tb*

**Table SI:** List of gRNA sequences used for gene knock-out/in and RNA interference.

**Table S2:** List of primers for gene knocked out and qPCR analysis.

**Table S3:** List of gRNAs synthesized for genome wide CRISPRi.

**Table S4:** List of differentially distributed targets sequences during *in vitro* screening of *M.tb*.

**Table S5:** List of *M.tb* drug target genes. The top 31 genes from the list show least similarity with the probiotics proteome.

**Table S6:** List of differentially distributed targets genes of *M.tb* growth inside Thp-1 derived macrophages.

